# Structural and Evolutionary Divergence of the RAG1/1L N-Terminal Zinc-Finger Domain

**DOI:** 10.64898/2026.01.24.700457

**Authors:** Junye Hong, Zhijun Liu, Eliza Martin, Kunyu Wei, David G. Schatz, Yuhang Zhang

## Abstract

RAG1/2 catalyzes V(D)J recombination to assemble antigen receptor genes, yet the zinc-finger element at the RAG1 N terminus has remained structurally and functionally uncharacterized. Using NMR spectroscopy, we determined the structure of the RAG1 N-terminal zinc-finger domain (NZD; residues 89–223). NZD adopts a compact, zinc-dependent fold comprising four α-helices organized into two interdigitated zinc-finger modules, ZFa and ZFb. Structural similarity searches across protein-structure databases revealed no solved homologs, establishing NZD as a previously undescribed zinc-finger fold. Comparative analyses show that NZD is broadly conserved across RAG1 and RAG1-like (RAG1L) proteins but has undergone lineage-specific remodeling, including the jawed vertebrate–specific acquisition of an additional α-helix (H2). Guided by structure prediction, we identified two NZD-like types in Chapaev transposases; one is highly similar to the RAG1L NZD, supporting an evolutionary link between RAG1/RAG1L and Chapaev NZDs. Similarity between ZFa and the single-zinc ZAD further suggests that NZD originated from an ancestral single-zinc fold. Together, these findings provide a structural framework for mechanistic and evolutionary analyses of RAG1.

## INTRODUCTION

The recombination-activating gene 1 (RAG1) and its cofactor RAG2 initiate the assembly of immunoglobulin and T cell receptor genes through V(D)J recombination.^1,2^ The RAG complex is a heterotetramer composed of two RAG1 and two RAG2 subunits. Within this complex, RAG1 serves as the catalytic subunit that recognizes and cleaves the recombination signal sequence (RSS) to trigger recombination, whereas RAG2 functions as a regulatory and structural cofactor.^1–4^ In mouse RAG1, the polypeptide can be divided into an N-terminal noncatalytic region (residues 1–384) and a C-terminal catalytic core.^5–9^ The latter mediates RSS recognition and DNA cleavage and has been extensively characterized by structural and biochemical studies. By contrast, the N-terminal region (residues 1–384) remains poorly understood at both functional and structural levels, especially the segment spanning residues 89–215, which has long been predicted and experimentally supported by biochemical data to form a “putative zinc finger”.^10–12^ This interval lacks experimental structural information as well as direct mechanistic evidence linking it to complex assembly, RSS selectivity, or nuclear localization.

Genetic studies and mouse models indicate that the N terminus is essential for immune homeostasis in vivo.^11,13–18^ Diverse N-terminal deletions or truncations cause severe immunodeficiency,^18–20^ arrested lymphocyte development,^13^ delayed humoral responses,^21^ and altered V(D)J repertoires.^11,13^ Although this region does not directly participate in DNA catalysis, it likely modulates V(D)J recombination. Prior work suggests that in the RAG1 N terminal region, the extreme N terminus (residues 1–88) is largely disordered;^10^ residues 89–215 are predicted by secondary-structure and coordination analyses to constitute zinc-finger domains;^10,12,17^ the next segment extending to residue 256 contains a positively charged, nuclear localization signal (NLS)-like/nucleolar localization signal (NoLS) motif;^22–24^ and residues 256–384 form an E3 ubiquitin ligase domain—the only N-terminal module that has, to date, been structurally resolved (**Figure 1A**).^25,26^

**Figure 1.**
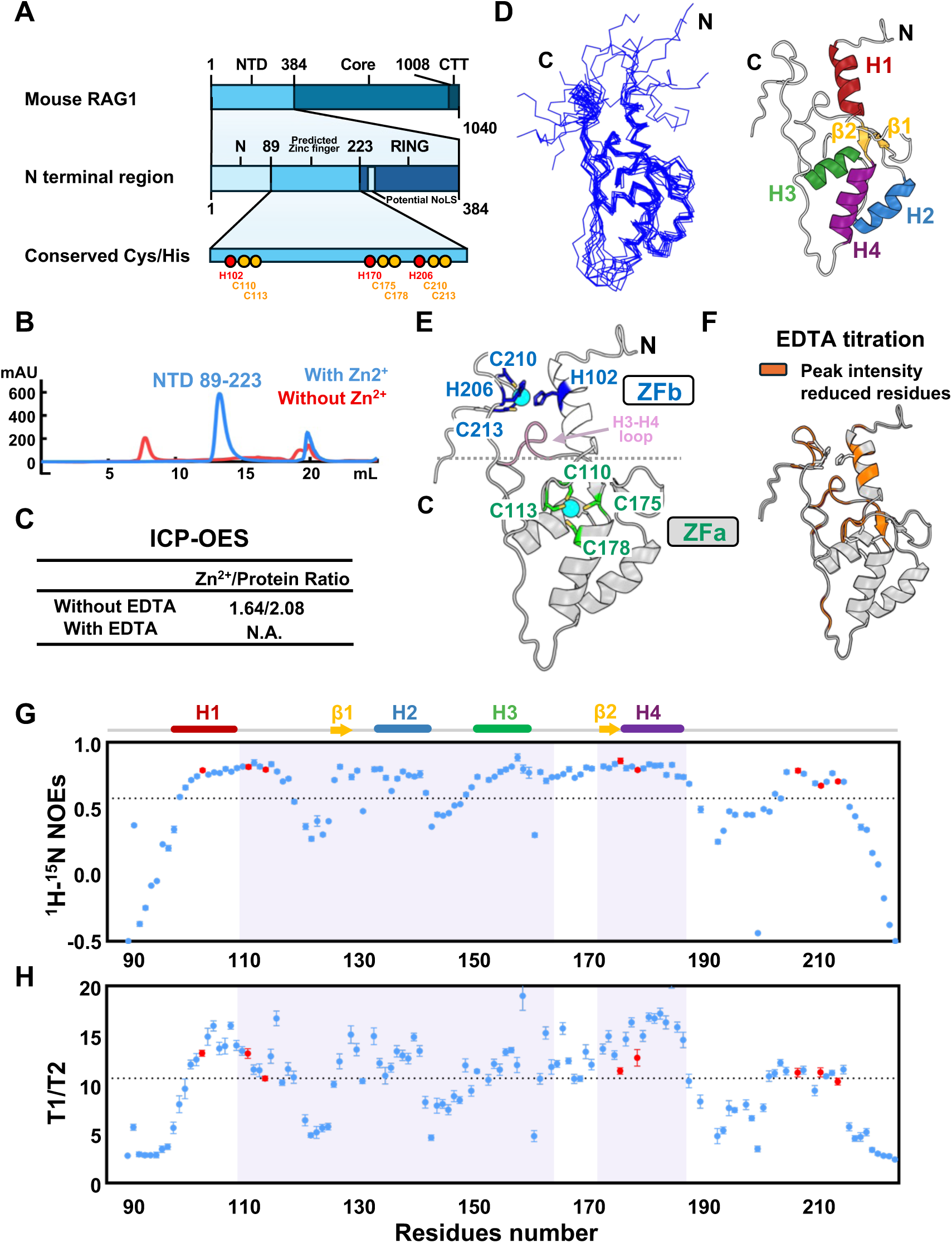
Structural analysis of the RAG1 NZD. (**A**) Domain architecture of mouse RAG1, showing the disordered N-terminal region (residues 1–88), the NZD (residues 89–223), a nucleolar localization signal (NoLS) motif, and the RING (Really Interesting New Gene) E3 ligase domain (residues 261-384). The intact N-terminal region and NZD are shown at increasing levels of magnification; putative Zn²⁺-coordinating residues in the NZD are marked as red dots (His) and yellow dots (Cys). NTD, N-terminal domain; CTT, C-terminal tail. (**B**) Size-exclusion chromatography (SEC) profiles of NZD under Zn²⁺-coordinated (blue trace) and Zn²⁺-depleted (red trace) conditions. The zinc-bound protein elutes as a monomer peak, whereas EDTA chelation yields an earlier-eluting oligomeric peak. (**C**) ICP-OES quantification of Zn²⁺ stoichiometry indicates two Zn²⁺ ions per NZD monomer. The figure shows the results from two independent measurements. (**D**) Left: ensemble of the 10 lowest-energy NMR models (ribbon representation). Right: NZD structure (cartoon representation) comprising four α-helices (H1–H4) connected by loops. Helices H1, H2, H3, and H4 are colored red, blue, green, and purple, respectively. (**E**) Cartoon representation of the zinc-binding architecture of NZD, which comprises two interdigitated zinc fingers, ZFa (dark grey) and ZFb (light grey). The zinc coordinated residues in ZFa (green) and ZFb (blue) are shown as sticks. Both Zn^2+^ are shown in cyan spheres, and the ZFb loop between helices H3 and H4 is highlighted in pink. (**F**) Mapping of residues exhibiting resonance intensity loss upon EDTA titration in NMR experiments. Residues showing >60% peak-intensity reduction at a 1:2 (protein:EDTA) molar ratio are highlighted in orange on the structure. (**G**) ¹H–¹⁵N steady-state heteronuclear NOE of NZD. NOE values were calculated as the ratio of peak intensities recorded with and without ¹H saturation (I_sat/I_ref). Error bars were obtained by propagating the spectral noise (SD) from the saturated and unsaturated spectra. The ZFa region is indicated by light-purple shading, and Zn²⁺-coordinating residues are marked in red. (**H**) T₁/T₂ ratios of NZD. T₁ and T₂ values were obtained by fitting peak intensities to single-exponential decays, and T₁/T₂ ratios were then calculated. Error bars represent fitting uncertainties in T₁ and T₂ propagated to the ratio. The ZFa region is indicated by light-purple shading, as in panel (G).

From an evolutionary perspective, RAG and its ancestral RAG-like transposases (RAG1L/RAG2L) provide a framework for explaining how a mobile genetic element was domesticated to execute site-specific immune recombination.^1,27–29^ Although the catalytic cores of RAG1 and RAG1L are relatively conserved, their N-terminal regions exhibit far greater sequence divergence,^28,30,31^ implying that the N terminus constitutes a hotspot for host adaptation and functional diversification. A key question is whether early invertebrate RAG1L proteins carried a folded N-terminal domain corresponding to the mouse aa 89–215 segment, whether its copy number and arrangement varied, and what evolutionary mode it followed. Answering these questions will not only illuminate the origin of the RAG1 N terminus but also clarify how RAG transitioned from a transposase-like enzyme to a regulated recombinase with precise substrate selectivity. Thus, the uncharacterized aa 89–215 “zinc-finger candidate” represents a missing link for understanding RAG1 regulation, as modeling its mechanistic role and tracing its evolutionary trajectory across vertebrate and invertebrate lineages remain difficult without an atomic-level structure.

To address these issues at the structural level, we focused on the 89–215 segment of mouse RAG1 (experimentally extended to 89–223 to ensure stable folding) and determined its three-dimensional structure by nuclear magnetic resonance (NMR) spectroscopy under zinc-bound conditions. We examined the role of metal coordination in folding and oligomerization and quantified the Zn²⁺:protein stoichiometry by inductively coupled plasma optical emission spectrometry (ICP-OES), revealing a 2:1 ratio.^32–34^ This result is consistent with prior experimental inferences.^10^ Combining cross-species structural searches and similarity analyses in Foldseek^35,36^ as well as AlphaFold3 (AF3) predictions,^37,38^ we observed a gradual transition from a single-finger to a dual-finger architecture accompanied by a redistribution of surface electrostatics. These results define a previously uncharacterized double zinc-finger fold, the RAG1 N-terminal zinc-finger domain (NZD), in which both metal-binding sites are essential for domain stability; disruption of either site markedly reduces recombination efficiency. Together, structural, biochemical, and evolutionary evidence supports the view that this zinc-finger domain is a co-evolved regulatory module that may have helped endow RAG1 with the selectivity and programmability required for its evolutionary transition.

## RESULTS

### The RAG1 NZD coordinates two zinc ions

Secondary-structure predictions indicate that the RAG1 N terminus contains multiple potential zinc-coordinating motifs. In addition to the RING E3 ligase domain (residues 256–384), a second hotspot of putative zinc binding centers on residues 89–215 (**Figures 1A and S1**). On the basis of this interval as a starting construct, we systematically evaluated C-terminal extensions and identified residues 89–223 as the optimal expression boundary. This span encompasses all predicted Zn²⁺-binding sites and yields a stable, soluble protein. Expressed in M9 minimal medium supplemented with Zn²⁺ and purified after tag removal, the construct eluted as a single peak by size-exclusion chromatography (SEC) (**Figure 1B**). The elution volume matched that expected for a monomer, indicating proper folding, monodispersity, and suitability for high-resolution NMR data collection. Sequence analysis of residues 89–223, which contains multiple CXXC motifs, suggests more than one metal-binding module (**Figure S1**). ICP-OES quantified two Zn²⁺ ions per protein molecule (**Figure 1C**). This is consistent with previously reported ICP-MS data for the RAG1 87–217 fragment,^10^ indicating that the stable protein domain we obtained contains two zinc binding modules. By contrast, expression under zinc-deficient conditions and purification in the presence of EDTA shifted the main SEC peak toward a higher apparent molecular weight and produced a void-volume peak (**Figure 1B**), consistent with hydrophobic exposure and nonspecific oligomerization—a hallmark of zinc-finger domains from which the metal ions have been removed. We therefore conclude that a bimetallic coordination scaffold is required to maintain the fold. As this domain is the most N-terminal zinc binding region of RAG1, we termed it NZD and proceeded to determine its solution structure so as to define its topology and coordination geometry at atomic resolution.

### NMR structure of the RAG1 NZD

We determined the NZD structure by high-resolution NMR spectroscopy. The ^15^N-labeled sample exhibited a well-dispersed ^1^H–^15^N HSQC spectrum (**Figure S2A**). Using a ^13^C,^15^N labeled sample, we acquired three-dimensional spectra and completed backbone and side-chain assignments. Structure calculations incorporating NOE-derived distance restraints, dihedral-angle restraints, and hydrogen-bond restraints resulted in a well-defined ensemble; the ten lowest-energy conformations superimpose with a backbone RMSD (Root-mean-square deviation) of 2.0 Å (**Figure 1D , left**).

The fold is composed of four α-helices (H1–H4) connected by intervening loops and two β-strands, which together assemble into two integrated zinc-finger modules (**Figure 1D , right and 1E**). The lower module, ZFa, is a C4-type zinc finger primarily formed by helices H2–H4 and constitutes the folding core of ZFa. The zinc-chelating site, coordinated by C110, C113, C175, and C178, is positioned above the helical core (**Figures 1E and S2B**). In contrast to ZFa, ZFb is a C2H2-type zinc finger involving the N-terminal helix (H1), the C-terminal segment, and part of the H3–H4 loop, and is coordinated by H102, H206, C210, and C213 (**Figures 1E and S2B**). The two zinc-finger modules interlock through loop–loop interactions to form an integrated fold. To assess the contribution of zinc coordination to NZD stability, we performed EDTA titration experiments. In the ^1^H–^15^N HSQC spectra, signals from coordinating residues decreased in intensity and exhibited slow-exchange behavior (**Figures S3A–S3C**). In ZFa, EDTA-induced signal attenuation was predominantly localized to the zinc-coordinating region (**Figure 1F**), indicating that the ZFa zinc-finger module is stabilized by the combined effects of helical packing and zinc-finger coordination. By contrast, ZFb exhibited extensive signal attenuation throughout the module, indicating that zinc coordination is essential for maintaining its conformation.

To further characterize the dynamics of the two zinc-finger modules, ^15^N heteronuclear NOE experiments were performed to identify fast protein backbone motion. The values differ markedly between ZFa and ZFb (**Figure 1G**), with average values of 0.65 ± 0.02 and 0.44 ± 0.07, respectively, indicating distinct backbone dynamic properties. Consistently, in the ^15^N relaxation experiments, the T1/T2 ratios also show differences between ZFa (11.3 ± 0.4) and ZFb (8.8 ± 0.7) (**Figure 1H**). ZFb exhibits a substantially lower average ratio, indicative of enhanced backbone mobility. Taken together, these results indicate that, compared with the more rigid and tightly packed ZFa module, ZFb adopts a more dynamic and loosely packed conformation, revealing substantial heterogeneity in their structural features.

### NZD defines a novel protein fold

To determine whether NZD represents a unique protein fold, we searched structural databases using Foldseek with the TM-align structural alignment algorithm, which reports a TM-score as a measure of global fold similarity.^36,39^ No matches were identified in the experimental structure database (**Figure 2A**), supporting the conclusion that NZD is a previously uncharacterized zinc-binding fold. Searches against predicted-structure databases revealed that the majority of alignment hits (63.3%) are annotated as RAG1 or RAG1-like proteins (**Figure 2B**). More than 90% of these hits correspond to vertebrate RAG1, with only a small number scattered among invertebrate lineages. Notably, this subset of predicted structures consistently exhibits a dual zinc-finger organization, characterized by spatially proximal Cys and/or His residues capable of coordinating zinc ions. Although a small fraction of the hits were annotated as zinc-finger associated domain (ZAD) (7.9%) or histidine–asparagine–histidine domain (HNH) containing proteins (5.9%), both of these domain types harbor only a single zinc-binding site and lack the dual zinc-finger module architecture observed for NZD.^40–42^ Therefore, we conclude that NZD represents a previously unclassified, novel zinc-finger fold.

**Figure 2.**
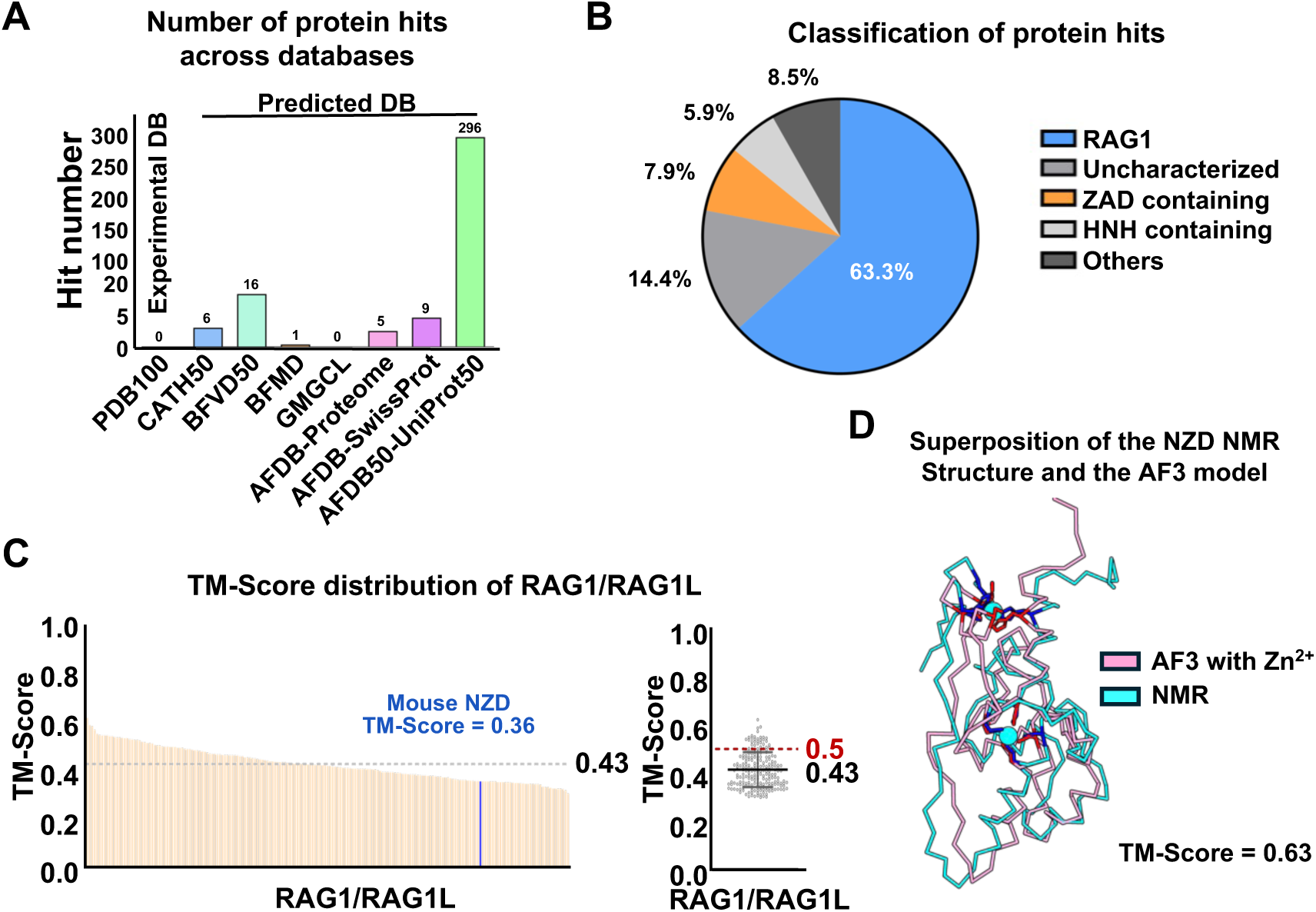
NZD adopts a previously uncharacterized zinc-finger fold. **(A)** Foldseek structural similarity searches (TM-align mode) using the NZD structure as the query did not identify any experimentally determined structural homologs in the databases surveyed. PDB100 (non-redundant experimental PDB structures), CATH50, BFVD, BFMD, GMGCL, AFDB-SwissProt, AFDB-Proteome, and AFDB-UniProt50. The numbers above each bar indicate the number of hits in each database. **(B)** Classification of protein hits identified in (A). Most hits mapped to RAG1/RAG1L homologs, with smaller fractions annotated as uncharacterized proteins, ZAD containing proteins, HNH-domain proteins, or other categories. Categories: RAG1, jawed vertebrate RAG1 and invertebrate RAG1L; ZAD containing, proteins containing a zinc-finger–associated domain (ZAD); HNH containing, proteins containing an HNH (His–Asn–His) domain; Uncharacterized, proteins lacking functional annotation; Others, all remaining hits. **(C)** Left: histogram of TM-score for RAG1/RAG1L hits identified in (A). The gray dashed line indicates the mean TM-score across all hits; the mouse RAG1 NZD is highlighted in blue. Right: scatter plot of TM-scores for all structural hits retrieved from predicted structure databases. The red dashed line marks TM-score = 0.5, a commonly used heuristic threshold for classifying two proteins as sharing the same fold. **(D)** Structural superposition of the NMR-determined NZD structure (cyan) with an AlphaFold3-predicted model of mouse RAG1 89–223 (pink), which was generated with Zn²⁺ ions explicitly included in the prediction. The resulting TM-score of 0.63 indicates good overall agreement in both global topology and Zn²⁺-coordination geometry.

Although vertebrate RAG1 proteins share relatively high sequence similarity, analysis of predicted structures indicates that, in the absence of Zn²⁺, NZD exhibits limited alignment quality, with an average TM-score of 0.43 across all hits. Consistent with this, the mouse RAG1 NZD model from the predicted-structure database yields a TM-score of only 0.36 in this comparison (**Figure 2C**), suggesting that accurate prediction of NZD requires additional explicit ion-coordination constraints. To address this limitation, we used the sequence of mouse RAG1 NZD (residues 89–223), rather than the full-length protein, as the prediction target and performed independent structure prediction using AlphaFold3 (AF3). When zinc ions were explicitly included during prediction, agreement with the experimentally determined structure improved substantially (TM-score 0.63; RMSD 3.0 Å). Despite minor discrepancies, the predicted model closely recapitulates the overall fold and spatial organization of the experimental structure (**Figure 2D**). These results indicate that zinc-informed AF3 predictions of truncated NZD constructs provide a reliable structural framework for investigating NZD family homologs.

### Divergence of NZD architectures across RAG1/RAG1L lineages

NZD is a previously unrecognized, RAG1-specific zinc-finger domain that retains detectable sequence similarity across RAG1/RAG1L clades, providing a basis for exploring structural evolution. Using AF3 with explicit zinc-coordination priors, we predicted structures for representative RAG1 and RAG1L proteins (**Figure S4**). In jawed vertebrates, the predicted NZD is in close agreement with our NMR structure (**Figures 3A and S4**), consistent with strong sequence conservation. In contrast, comparative analyses of invertebrate RAG1L NZD indicate that in several early-branching metazoans (e.g., cnidarians, planarians), a subset of family-B and family-D RAG1L proteins carries only a single ZFa-like N-terminal zinc-finger module (**Figures 3A and S4**), whereas other RAG1L proteins—either within the same species or in other invertebrates—clearly contain a second zinc-finger module (ZFb), as observed in mouse NZD (**Figures 3A and S4**). Consistent with this, in vivo recombination assays show that mutating zinc-coordinating residues in either ZFa or ZFb markedly reduces RAG1-dependent recombination but does not completely abolish it (**Figure 3B**).

**Figure 3.**
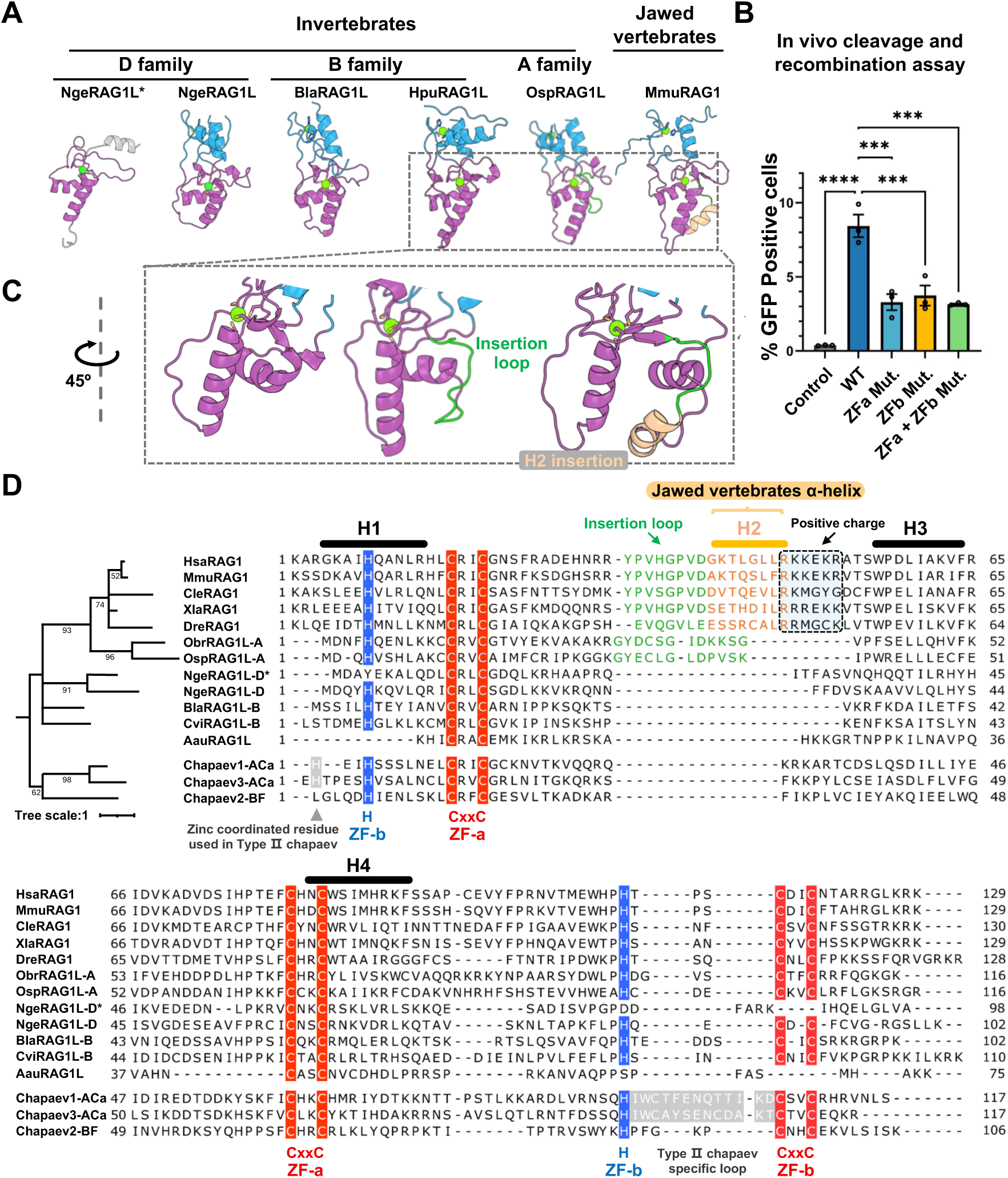
Evolutionary diversification of the NZD fold across RAG1/RAG1L. **(A)** Representative AlphaFold3-predicted structural models, alongside the experimentally determined NMR structure of mouse RAG1 NZD, reveal the architectural diversity of NZD across evolution. In invertebrate RAG1L proteins, most members of families A, B, and D adopt a dual zinc-finger architecture comprising ZFa and ZFb (see also Fig. S4). Notably, family A features an insertion loop (light green) between helices H1 and H3 (see also panel C). * indicates that the NgeRAG1 NZD (referred to here) lacks ZFb.. In jawed vertebrate RAG1, an additional α-helix (H2, orange) appears between H1 and H3, forming part of the ZFa module. ZFa (purple), ZFb (blue), and Zn²⁺ (green spheres). **(B)** Functional validation of zinc-finger requirements for RAG1 activity using GFP reporter recombination assays in Expi293 cells. Cells were transfected with the p290GFP reporter together with expression constructs for full-length wild-type or mutant RAG1 (with Zn²⁺-coordinating residues mutated) and RAG2. GFP-positive cells indicate successful recombination. Both ZFa and ZFb are required for robust RAG1 function; mutating any Zn²⁺-coordinating residue in either finger markedly reduces RAG1-mediated cleavage and recombination. Data are presented as mean ± s.e.m. from at least three independent experiments. Statistical significance was assessed by one-way ANOVA followed by Dunnett’s multiple-comparisons test (each mutant vs. wild-type RAG1). ***P < 0.001; ****P < 0.0001; ns, not significant. **(C)** Enlarged views of selected NZD structures from panel A, rotated by 45°, highlighting structural variation in protein fold arrangement. **(D)** Maximum-likelihood phylogeny of NZD sequences from the RAG1/RAG1L and Chapa domain sequences from Chapaev transposase families, inferred using IQ-TREE (see Materials and Methods), alongside the corresponding multiple sequence alignment. In the alignment, conserved Zn²⁺-coordinating residues are highlighted—histidines (His) in blue and cysteines (Cys) in red. Secondary structure elements are annotated above the alignment: α-helices H1, H3, and H4 are indicated by black bars; a family A–specific insertion loop between H1 and H3 is shown in green; the vertebrate-specific H2 helix within ZFa is marked in light orange, with a dashed box highlighting an additional basic residue–rich insertion. Gray triangles indicate type II Chapa domain–specific Zn²⁺-coordinating residues; gray shading marks the corresponding type II Chapa domain–specific loop.

Most RAG1/RAG1L homologs retain both ZFa and ZFb; however, RAG1L families A, B, and D commonly lack helix H2. In the RAG1L family-A, which most closely resembles jawed vertebrate RAG1 (**Figure S4**), an insertion appears in NZD between H1 and H3 (**Figure 3C**); based on AF3-predicted structures, this insertion does not form an α-helix but extends downward to partially cover the H4 surface that is exposed in families B and D (**Figures 3C and 3D**). In jawed vertebrates, while retaining the ancestral ZFa+ZFb framework, the extended family A sequence further elongates and folds into a new α-helical element (H2), followed by a positively charged loop. This basic surface formed by the loop is highly conserved (**Figures 3C and 3D**) and shows a stepwise increase in basic residues from cartilaginous fishes to bony fishes to amphibians, after which it stabilizes—suggesting a vertebrate-specific function subject to positive selection (**Figure 3D**). Conversely, jawed vertebrate NZDs display a newly emerged electropositive surface centered at the C terminus of helix H2 and the adjacent loop at the base of ZFa, which expands progressively across vertebrate lineages (**Figure S5**). These surface-potential shifts, likely driven by secondary-structure insertions, provide a plausible biophysical basis for potential functional expansion.

### Evolutionary origin of the RAG1 NZD

As a newly defined zinc-binding fold, the evolutionary origin of NZD remains unclear. Given that Transib transposases are the closest known relatives of RAG1/RAG1L, we surveyed reported Transib families and analyzed the organization of their N-terminal regions.^28,43,44^ All families encode a part or the whole canonical catalytic core domain and possess additional extensions relative to the catalytic core; however, structure prediction indicates that none of these extensions contains an NZD-like domain (**Figures 4A and S6**). Together with prior sequence-based analyses, these observations support the conclusion that NZD is distinct from the core and was not directly inherited from a common RAG1/Transib ancestor.

**Figure 4.**
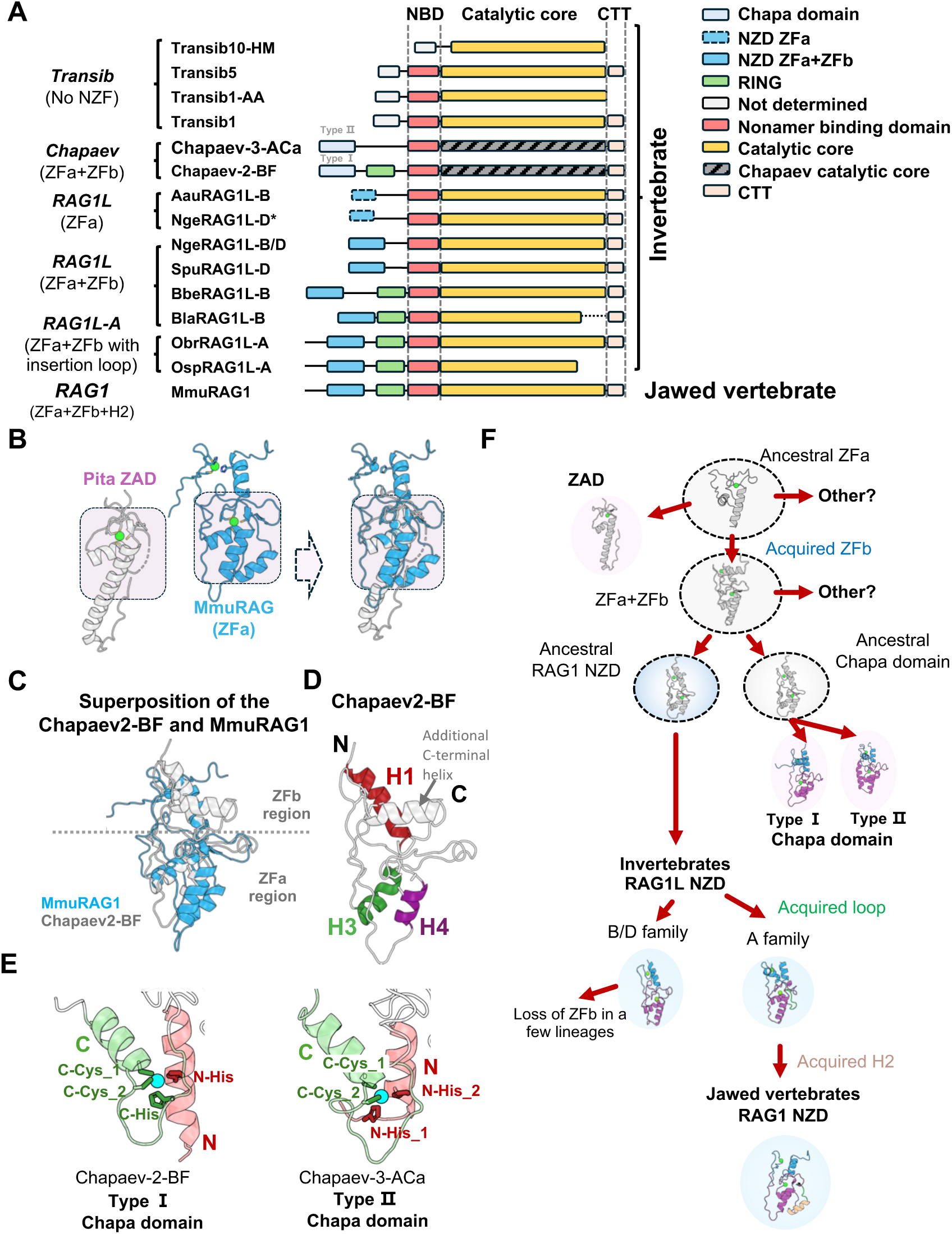
Evolutionary origin and trajectory of the RAG1 NZD. **(A)** Schematic comparison of domain architectures among the RAG1 recombinase, RAG1L, and Transib and Chapaev transposase families. No NZD-like domain was detected in Transib transposases. By contrast, Chapaev transposases contain an N-terminal NZD-like Chapa domain but show lower catalytic core sequence similarity to RAG1/RAG1L. Two Chapa domain subtypes (type I and type II) identified in Chapaev transposases are indicated. NBD, nonamer binding domain in jawed vertebrate RAG1 or NBD-like in invertebrate RAG1L; CTT, C terminal tail in jawed vertebrate RAG1 or CTT-like in invertebrate RAG1L. **(B)** Structural comparison of the ZAD of Drosophila melanogaster Pita (left; PDB: 7POK) and mouse RAG1 NZD (center), with the superposition shown on the right. The purple shaded region highlights the major overlap, primarily involving the ZFa of NZD, indicating local structural similarity. **(C)** Structural superposition of the AlphaFold3-predicted Chapaev-2-BF Chapa domain (gray) with the mouse RAG1 NZD structure (light blue) reveals a conserved ZFa/ZFb arrangement and overall NZD architecture. The gray dashed line indicates the boundary between the ZFa and ZFb zinc-finger modules. **(D)** The Chapa domain contains helices H1, H3, and H4 that correspond to those in mouse RAG1 NZD and includes an additional predicted C-terminal helix. However, as in invertebrate RAG1L NZD, it lacks the H2 helix characteristic of jawed-vertebrate RAG1. **(E)** Close-up views illustrating two distinct zinc-coordination patterns in the ZFb module of Chapa domains. In type I, zinc coordination involves one N-terminal residue and three C-terminal residues. In type II, zinc coordination involves two N-terminal residues and two C-terminal residues. The N-terminal and C-terminal portions of ZFb are shown as semi-transparent cartoon representations in light red and light green, respectively. Zinc-coordinating residues are depicted as sticks. **(F)** Proposed evolutionary trajectory of the RAG1 NZD. The dashed circle indicates a hypothetical evolutionary intermediate.

In the Foldseek-based structural search, we noted that although ZAD-containing proteins do not exhibit the overall fold characteristic of NZD, structural superposition revealed a degree of local architectural similarity between ZAD and the ZFa module of RAG1L NZD. Specifically, both contain helices H3 and H4 and a zinc-coordinated core, with approximately 23% sequence identity in this region (**Figures 4B , S7A, and S7B**), suggesting that they may have originated from a common or closely related zinc-finger ancestor. Despite this resemblance, ZAD and the ZFa module of NZD differ in several key features. First, unlike NZD, ZAD mediates dimerization.^40,45^ Second, ZAD lacks the prominent electropositive patch surrounding helix H4 that is characteristic of NZD ZFa (**Figure S7C)**. Together, these observations indicate that the similarity between ZAD and NZD ZFa is confined to a limited structural framework rather than reflecting a shared global fold.

The occurrence of the ZFb module appears to be evolutionarily contingent on the presence of ZFa and, from a structural perspective, ZFb is best interpreted as a derived elaboration of the ZFa scaffold rather than as an autonomous module. In Foldseek-based structural searches, only sporadic proteins, such as the Hypp6231 protein from *Branchiostoma lanceolatum* (**Supplementary Table 1**), yield predicted models that are compatible with the presence of a ZFb-like element. However, these proteins lack functional characterization and do not cluster into a coherent evolutionary group, precluding robust inference of their biological relevance. By contrast, available evidence indicates that Chapaev transposases, members of the CMC transposase superfamily, represent the only lineage in which N-terminal sequences (Chapa domain) with detectable similarity to RAG1L NZD are consistently retained.^44,46^ To examine this relationship, we predicted and analyzed the structures of N-terminal regions from multiple Chapaev transposases. These analyses revealed a compact zinc finger in Chapaev proteins that incorporates both ZFa and ZFb within a single, integrated fold (**Figure 4C**). Notably, the folds of Chapa domain and RAG1L NZDs are more similar to each other than either is to vertebrate RAG1 NZD, consistent with a shared ancestral architectural state. Both retain helices H1, H3, and H4 but lack helix H2 (**Figure 4D**), whereas H2 represents a later acquisition in the vertebrate lineage. In Chapa domain ZFb, zinc coordination is predicted to be polymorph**i**c, with at least two schemes: type I, one N-terminal plus three C-terminal ligands; or type II, two N-terminal plus two C-terminal ligands—the former mirroring the pattern in RAG1 NZD (**Figures 3C and 4E**). Relative to type I, type II carries an additional C-terminal insert that reassigns the zinc-chelating residues (**Figure 3D**). By comparison, ZFb ligands in RAG1 and RAG1L are highly conserved, typically comprising one His from the N terminus and one His plus two Cys from the C terminus as the type I Chapa domain (**Figure 3D**).

## DISCUSSION

We identify the RAG1 N-terminal zinc-finger domain as a previously unrecognized, zinc-finger fold specific to the RAG1 family. NMR together with ICP-OES demonstrates that NZD comprises two interdigitated zinc-finger modules, ZFa and ZFb, and that both metal-coordination sites are indispensable for structural stability and for efficient RSS cleavage and recombination in cells. No close match for NZD was found among solved structures, and high-scoring Foldseek hits map largely to vertebrate RAG1 N termini, underscoring family specificity. EDTA titrations further highlight the centrality of metal coordination: removing zinc collapses the fold, whereas zinc-informed, truncated domain predictions better recapitulate the experimental structure.

Structural comparative analyses across species and families reveal a conserved overall topology accompanied by lineage-specific remodeling of surface electrostatics. In invertebrate RAG1L, the H2 helix is absent; in family A, an insertion appears on one side of the H4 helix; and in jawed vertebrates, the H2 helix becomes a stable element within ZFa, with a conserved and progressively enlarged basic surface emerging at the ZFa/H2 base. Together with the strict requirement for bimetallic coordination, these features suggest that NZD functions primarily as a regulatory scaffold that fine-tunes protein–DNA and/or protein–protein interactions rather than serving as a catalytic moiety per se. The physiological role of NZD remains to be defined, and the vertebrate emergence of H2 and its lineage-specific electrostatic reconfiguration warrant targeted functional tests.

On the basis of these data, we propose an evolutionary model (**Figure 4F**) in which NZD originated from a distant, ZAD-related single-zinc-finger ancestor (ZFa-like domain) and subsequently acquired a second zinc finger through N- and C-terminal extensions. This two-zinc-finger (ZFa+ZFb) fold was then incorporated into the ancestors of both RAG1 and Chapaev transposases, where it gave rise to the ancestral NZD and the ancestral Chapa domain, respectively. The Chapa domain later diversified into two different fold types (Type I and Type II) on ZFb. The ancestral fold type of RAG1 NZD is more closely related to the Type I Chapa domain and continued to evolve within the RAG1/RAG1L lineage (**Figures S7A and 3D**). Starting from the A family RAG1L, an insertion loop appeared between helices H1 and H3; in jawed vertebrates, this loop further evolved into the helix H2, expanding a new basic surface at the base of ZFa (**Figure S5**). Collectively, these changes remodeled NZD into an evolutionarily coupled structural module, likely supporting the physiological transition of RAG1 function in vivo.

### Limitations of the study

Several limitations of this study should be noted. First, although we determined the solution structure of mouse RAG1 NZD and defined its folding topology, how NZD engages specific protein or DNA partners in the context of the intact RAG complex, and whether such interactions are modulated by the lineage-specific surface features identified here, remain to be elucidated through direct biochemical and structural analyses. Second, the evolutionary inferences presented here are largely based on structure-informed predictions and comparative analyses, while these approaches reveal coherent trends in NZD architecture and metal coordination across lineages, experimental validation of NZD or Chapa domain function in invertebrate RAG1L or Chapaev systems is currently lacking. Third, the evolutionary model proposed in this study is necessarily constrained by the data currently available, and may require refinement as additional invertebrate, and potentially viral, genomes and encoded proteins become accessible.

## ACKNOWLEDGMENTS

The authors thank Dr. Haixia Jiang from the Instrument Sharing Platform of the School of Life Sciences and Biotechnology at Shanghai Jiao Tong University for technical support with FACS experiments. This work was supported by the National Key R&D Program of China (2022YFE0132100, Y.Z.) and the National Natural Science Foundation of China (grant 32171250, Y.Z.).

## AUTHOR CONTRIBUTIONS

J.H., Y.Z., and Z.L. designed the study. J.H. and K.W. prepared protein and DNA constructs. J.H. and E.M. performed sequence alignment and phylogenetic analyses. J.H. and K.W. conducted FACS experiments. J.H. and Z.L. performed NMR experiments; Z.L. acquired and processed the NMR spectra, while J.H. carried out chemical shift assignments and structure calculations. J.H. conducted structural predictions and homologous structure searches. J.H. prepared the figures. Y.Z. wrote the manuscript. Y.Z. and D.G.S. revised the manuscript. All authors reviewed the results and approved the final version of the manuscript.

## SUPPLEMENTAL INFORMATION

Supplementary data including Supplementary Figure S1–S7.

## DATA AVAILABLE

The NMR structure of mouse NZD has been deposited in the Protein Data Bank (21WB). The AlphaFold3 predicted NZD structures used in Figures 3A and 4E have been deposited in ModelArchive under the following IDs: ma-wa0gi (BlaRAG1L NZD), ma-lctn2 (HpuRG1L NZD), ma-fjqnc (NgeRAG1L NZD), ma-qx8h7 (NgeRAG1L*), ma-69xz6 (OspRAG1L NZD), ma-dzh7q (Chapaev-2-BF Chapa domain) and ma-7ecx2 (Chapaev-3Aca Chapa domain) (https://modelarchive.org). All AlphaFold3 prediction data generated in this study have been deposited in Zenodo open-access online repository (https://zenodo.org/records/18073344).

## Methods

### Plasmid generation

The mRAG1 NZD (residues 89–223) used for protein purification and NMR spectroscopy was cloned into the pGEX-6P-1 vector, generating an N-terminal GST fusion followed by a PreScission protease cleavage site. For *in vivo* recombination assays, full-length mouse RAG1 was cloned into a pEBB mammalian expression vector with an N-terminal HA tag. Core RAG2 (residues 1–387) was cloned into a pTT5 mammalian expression vector with an N-terminal maltose-binding protein (MBP) tag. The RAG1 zinc-finger mutants, ZFa (C110A/C113A) and ZFb (C210A/C213A), were generated by site-directed mutagenesis of the pEBB-HA-RAG1 plasmid.

### Protein expression and purification

GST-tagged mRAG1 NZD (residues 89–223) was expressed in *E. coli* BL21(DE3) grown in either LB or M9 medium, as required for specific experiments, and supplemented with 30 μM ZnSO₄. Protein expression was induced with 0.5 mM IPTG at an OD₆₀₀ of 0.6, and cultures were incubated overnight at 16 °C. For isotopically labeled samples, cells were cultured in M9 minimal medium containing 30 μM ZnSO₄ as well as ^15^N-labeled NH₄Cl and/or ¹³C-labeled glucose as the sole nitrogen and/or carbon sources.

Cells were harvested, resuspended in ice-cold lysis buffer (50 mM Tris-HCl pH 8.0, 500 mM NaCl, 25 μM ZnSO₄, 2 mM DTT), and lysed by high-pressure homogenization at 4 °C. The lysate was centrifuged at 18,000 × *g* for 40 min at 4 °C, and the GST fusion protein in the supernatant was purified on a glutathione agarose column. On-bead cleavage was performed to remove the tag using PreScission protease, and the cleaved RAG1 NZD was collected in the flow-through. The protein was subsequently purified by size-exclusion chromatography on a Superdex 75 Increase column with buffer containing 10 mM MOPS pH 6.5, 150 mM NaCl, and 0.2 mM TCEP. Peak fractions were pooled, concentrated, and either used immediately or flash-frozen at –80 °C.

### Data collection and structure determination

¹⁵N/¹³C-labeled mouse RAG1 NZD samples were prepared at a protein concentration of 0.6 mM in buffer (10 mM MOPS pH 6.5, 150 mM NaCl, 0.2 mM TCEP) containing 10% D₂O (v/v). Multidimensional NMR spectra, including 2D ¹H–¹⁵N HSQC and the triple-resonance experiments HNCACB, CBCA(CO)NH, HBHA(CO)NH, CC(CO)NH, HNCO and HN(CA)CO, were recorded at 25 °C on a Bruker AVANCE III HD 600 MHz spectrometer equipped with a TCI cryoprobe. The 3D ¹⁵N-NOESY–HSQC spectrum was acquired at 25 °C on a Bruker AVANCE III HD 900 MHz spectrometer.

NMR data were processed in NMRPipe using standard procedures.^47^ All spectra were analyzed in NMRFAM-Sparky^48^ for backbone and side-chain resonance assignments and NOE peak picking. NOE cross-peaks from ¹⁵N-resolved NOESY–HSQC spectra were converted into interproton distance restraints. Backbone dihedral angle restraints (φ and ψ) were derived from assigned backbone chemical shifts using TALOS-N,^49^ and hydrogen-bond restraints were incorporated for slowly exchanging backbone amide protons identified in hydrogen–deuterium exchange experiments.

Three-dimensional solution structures of RAG1 NZD were calculated in XPLOR-NIH^50^ using NOE-derived distance restraints, TALOS-N–based dihedral angle restraints, and hydrogen-bond restraints. Zinc ions were included during structure calculations in XPLOR-NIH by loading the zinc-finger topology and parameter set distributed with XPLOR-NIH (“*toppar/extra/zn-finger.top*” and “*toppar/extra/zn-finger.par*”). Zinc–ligand coordination was enforced using the corresponding zinc-finger patches and the associated bond/angle terms to maintain a tetrahedral coordination geometry during simulated annealing. A total of 100 structures were generated by simulated annealing from randomized starting conformations and iteratively refined until no significant NOE or dihedral-angle violations remained. The 10 lowest-energy conformers were selected to represent the final structural ensemble of RAG1 NZD. Structures were visualized and rendered in PyMOL.

### Backbone ^15^N relaxation and heteronuclear NOE

Backbone ^15^N relaxation measurements were performed on 15N-labeled RAG NZD at a protein concentration of 0.6 mM. Longitudinal relaxation (T1) data were collected using eight relaxation delays (2, 20, 40, 80, 160, 320, 640, and 1280 ms) with a 2 s recycle delay.

Transverse relaxation (T2) was measured using ten relaxation delays (0, 20, 40, 80, 100, 120, 160, 200, 300, and 400 ms). Peak intensities were extracted on a per-residue basis and fitted to single-exponential decays to obtain T1 and T2, and T1/T2 ratios were computed per residue. Steady-state ^1^H–^15^N heteronuclear NOEs were acquired as an interleaved pair of experiments recorded with and without 1H saturation. For the saturated experiment, spectra were recorded with a 2 s recycle delay followed by a 3 s 1H saturation period, whereas the reference experiment was acquired without saturation using a 5 s recycle delay. Heteronuclear NOE values were calculated as the intensity ratio between spectra acquired with and without ^1^H saturation (I_sat/I_ref). Uncertainties were estimated from spectral signal-to-noise and the fitting errors and propagated to derived quantities.^51^

### NMR titration experiment

NMR titration experiments were performed in buffer containing 10 mM MOPS pH 6.5, 150 mM NaCl, and 0.2 mM TCEP. ¹⁵N-labeled NZD (50 μM) was titrated with EDTA at protein:EDTA molar ratios of 1:0, 1:1, 1:2, and 1:5, and ¹H–¹⁵N HSQC spectra were acquired at each titration point. NMR data were processed as described in the “Data collection and structure determination” section, and peak intensities were normalized to those of the sample containing ¹⁵N-labeled NZD only (1:0).

### In vivo cleavage and recombination assay

Expi293 cells were co-transfected with the p290GFP reporter plasmid (1.5 μg), an MBP–RAG2 expression plasmid (1.5 μg), and an HA–RAG1 expression plasmid (1.5 μg) encoding either wild-type RAG1 or zinc-finger mutants, including ZFa (C110A/C113A), ZFb (C210A/C213A), or the combined ZFa+ZFb mutant (C110A/C113A/C210A/C213A). For negative controls, cells were transfected with p290GFP (1.5 μg) and an MBP control plasmid (3.0 μg). In all conditions, the total amount of DNA was adjusted to 4.5 μg per transfection. Forty-eight hours after transfection, cells were harvested, washed twice with ice-cold PBS containing 1% FBS, and stained with DAPI (4ʹ,6-diamidino-2-phenylindole) for 20 min at room temperature in PBS supplemented with 1% FBS prior to flow-cytometric analysis. Dead cells were excluded based on DAPI staining, and the fraction of GFP-positive events was quantified. The percentage of GFP-positive cells was used as a readout of RAG-dependent cleavage and recombination efficiency, as described previously.^52^

### Analytical size-exclusion chromatography

Size-exclusion chromatography (SEC) was performed on an NGC chromatography system (Bio-Rad Laboratories, Hercules, CA, USA) equipped with a Superdex 75 Increase 10/300 GL column (Cytiva, Marlborough, MA, USA) at 4 °C in buffer containing 10 mM MOPS pH 6.5, 150 mM NaCl, and 1 mM DTT. Target proteins were diluted to 30 µM, stored at 4 °C and then subjected to chromatography. For each run, 500 µL of protein sample was loaded onto the column, and elution was monitored by measuring absorbance at 280 nm. The oligomeric state of the protein was inferred from the retention time of the elution peak.

### Determination of NZD Zn²⁺-coordinating stoichiometry

To generate a Zn-depleted control, RAG1 NZD was expressed in M9 minimal medium without ZnSO₄ supplementation, using the same induction and purification procedures as for Zn-replete samples. Following purification, Zn-depleted RAG1 NZD was diluted to a protein concentration of 100 μM in the purification buffer. The sample was then incubated with 10 mM EDTA at 30 °C for 1 h to chelate residual Zn²⁺. After incubation, the sample was centrifuged at 17,000 × g for 5 min to remove any precipitates, and the supernatant was further purified by size-exclusion chromatography on a Superdex 75 Increase column equilibrated in the same buffer containing 1 mM EDTA.

Zinc content in Zn-replete and Zn-depleted protein samples was quantified by inductively coupled plasma–optical emission spectroscopy (ICP-OES) using an Avio 500 spectrometer (PerkinElmer), monitoring Zn emission at 206.2 nm. The Zn-to-protein molar ratio was calculated from the Zn concentration measured by ICP-OES and the protein concentration determined for each purified sample.

### Structural homology searches

Protein structural homology searches were performed using the Foldseek online server (https://search.foldseek.com; accessed October 24, 2025).^36^ As the query structure, we used the representative NMR structure of the mouse RAG1 NZD (model 9 from the NMR ensemble). The target databases comprised Foldseek prebuilt databases, AlphaFold/UniProt50 v6, AlphaFold/Swiss-Prot v6, AlphaFold/Proteome v6, BFVD 2023_02, CATH50 4.3.0, BFMD 20240623, and GMGCL 2204, as well as the experimentally determined structure database PDB100 20240101. Structural similarity searches were carried out with the “Foldseek-search” workflow in TM-align mode, in which Foldseek realigns candidate hits using the global TM-align algorithm and reports TM-scores as the primary ranking and statistical metric. All other parameters were kept at their default settings. Search results were exported in JSON format for downstream manual inspection and classification of hits into protein categories based on UniProt annotation (**Supplementary Table 1**).

### AlphaFold 3 structure prediction

Structures of NZDs from jawed vertebrate RAG1, invertebrate RAG1L, and Chapa domains from Chapaev transposases, as well as the N-terminal regions of Transib transposases, were predicted using the AlphaFold Server (AlphaFold 3; https://alphafoldserver.com).^37,38^ For each NZD and each Transib N-terminal region, separate prediction jobs were run specifying 0, 1, or 2 Zn²⁺ ions, with all other parameters kept at their default settings. The resulting models were visually inspected, and any models in which Zn²⁺ ions were not coordinated by the expected ligating residues or displayed unrealistic coordination geometries were discarded. For each NZD and each Transib N-terminal region, the top-ranked model (Model 0) from the predictions was selected as the reference model for subsequent analyses. All the sequences used are listed in Supplementary table 2.

### Sequence Alignment

Multiple sequence alignments (MSAs) of protein sequences were generated using ClustalX 2.1. Alignments were computed with the “Complete Alignment” option, using the BLOSUM30 substitution matrix, a gap opening penalty of 10, and a gap extension penalty of 0.2. Iterative refinement was enabled at each alignment step. When residues in the C-terminal region that are known to be locally conserved were not properly aligned, the corresponding region was refined using the “Realign Selected Range” function in ClustalX, followed by manual inspection. The final MSA was exported and visualized in Jalview 2.11 for interactive inspection, annotation, and figure preparation.

Pairwise percent identities between sequences were calculated from an MSA generated using the Clustal Omega multiple sequence alignment web server at EMBL-EBI (https://www.ebi.ac.uk/jdispatcher/msa/clustalo) with default settings, which employ seeded guide trees and HMM profile–profile techniques to generate accurate protein MSAs. The resulting percent-identity matrix was exported and used for downstream comparison and plotting.

### Phylogenetic Analysis

The sequences of NZD and Chapa domain were extracted from RAG1, RAG1L (Families A, B, and D), and Chapaev transposases and subjected to multiple sequence alignment for phylogenetic analysis. Model selection was performed based on the Akaike Information Criterion (AIC) and Bayesian Information Criterion (BIC), which identified LG+I+G4 (Le and Gascuel matrix with invariant sites and a discrete Gamma distribution with 4 categories) as the best-fit substitution model. A maximum likelihood (ML) tree was constructed using IQ-TREE^53^ with 1,000 ultrafast bootstrap replicates to assess branch support. To address phylogenetic uncertainty due to the short sequence length of the NZD domain, branches with bootstrap support values below 50% were collapsed. The resulting tree corresponds to the topology shown in Figure 3D.

## Supplemental information

**Figure S1.**
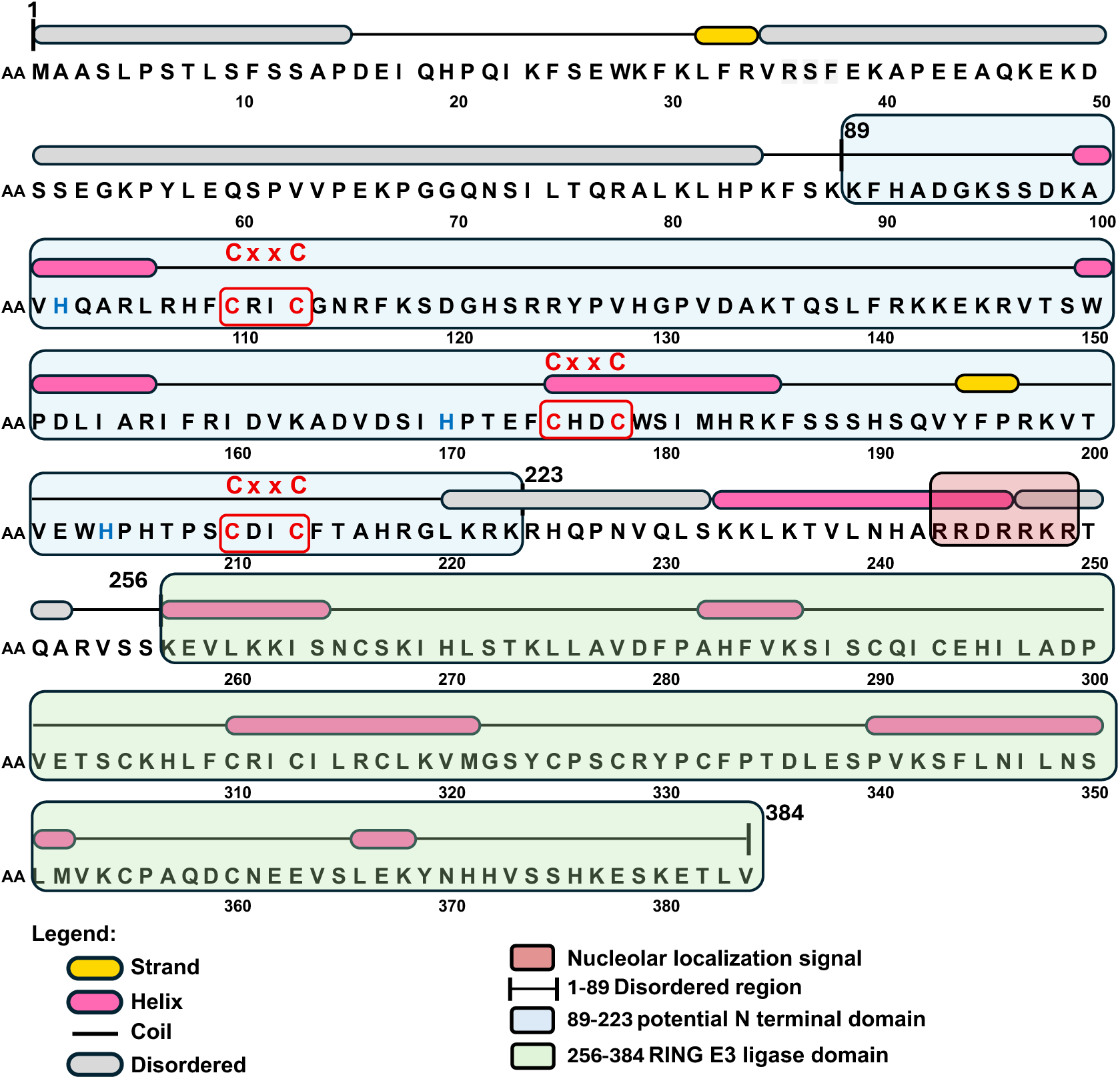
Secondary structure prediction of the mouse RAG1 N-terminal, related to Figure 1. Secondary structure prediction for the N-terminal region (residues 1–384) of mouse RAG1, generated using the PSIPRED. Different secondary-structure elements are indicated by distinct colored bars as shown in the figure. The NZD region (residues 89–223), E3 domain (residues 256–384), and positively charged residues corresponding to the nucleolar localization signal are highlighted with blue, green, and red shading, respectively.

**Figure S2.**
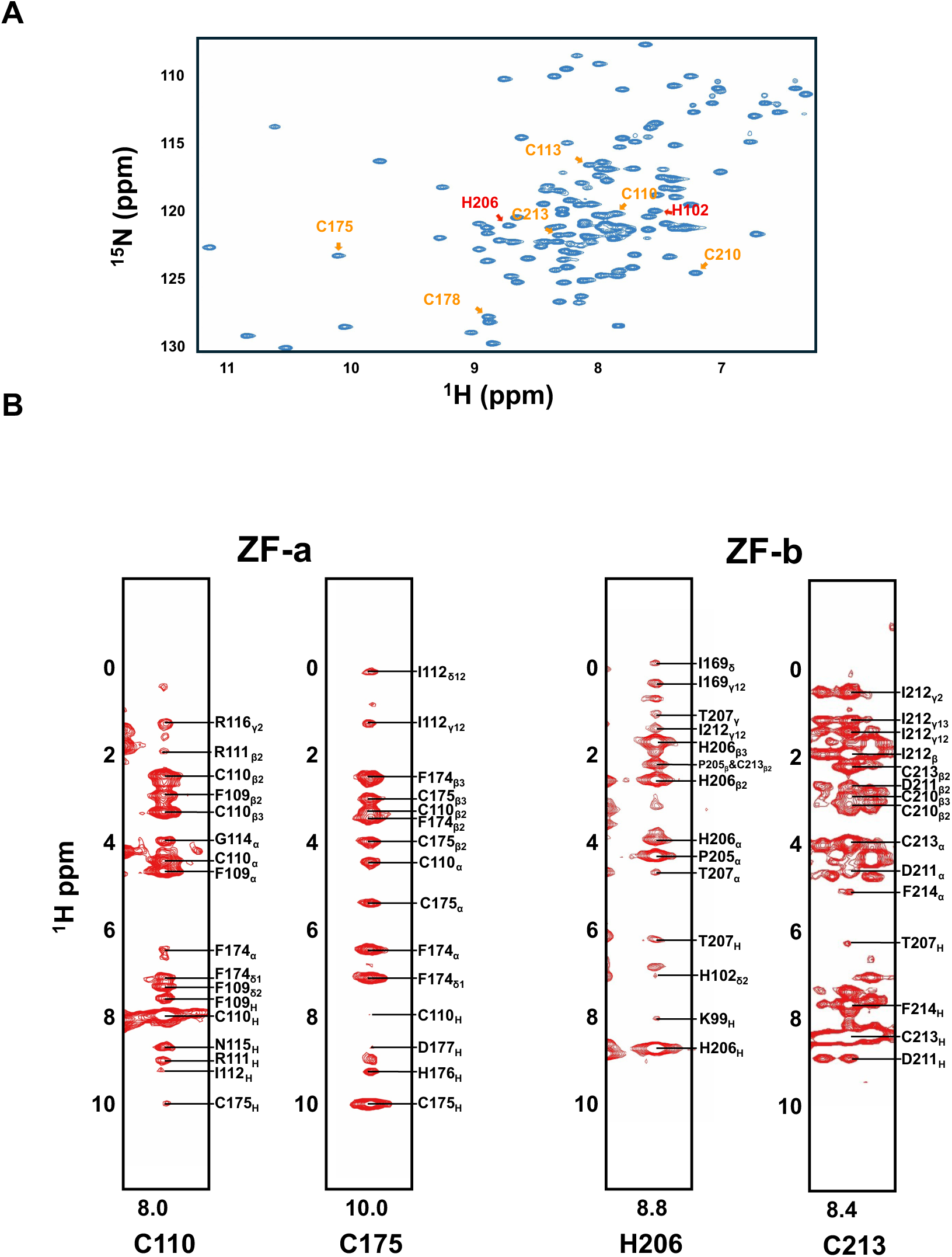
NMR resonance assignments of zinc-coordinating residues, related to Figure 1. **(A)** ¹H–¹⁵N HSQC spectrum of NZD under Zn²⁺-bound conditions, showing well-dispersed and sharp cross-peaks, indicative of a well-folded structure. Zn²⁺-coordinating residues are labeled in red (His) and yellow (Cys). **(B)** Selected representative NOE signals from the ¹⁵N-NOESY-HSQC spectrum, highlighting Zn²⁺-coordinating residues C110 and C175 in ZFa, and H206 and C213 in ZFb of NZD. Each strip is centered on the amide resonance of the labeled residue and shows intra- and inter-residue NOEs consistent with the close spatial proximity of residues around the Zn²⁺-binding sites.

**Figure S3.**
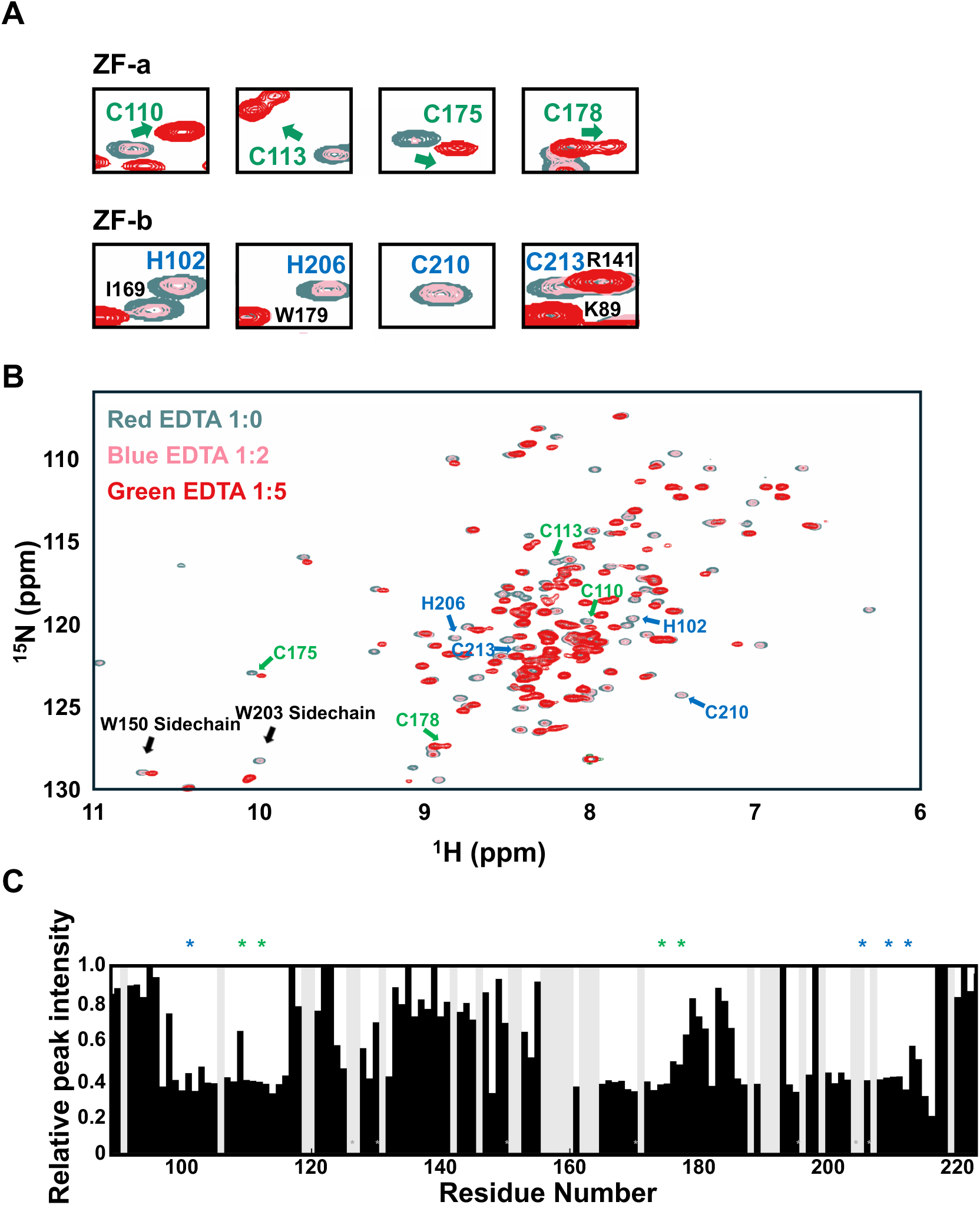
Loss of Zn²⁺ destabilizes NZD fold, related to Figure 1. **(A)** Local resonances of zinc-chelating residues in ^1^H–^15^N HSQC spectra, showing changes in peak intensity and chemical shift during EDTA titration at NZD:EDTA molar ratios of 1:0 (green), 1:2 (pink), and 1:5 (red). **(B)** Overlay of ^1^H–^15^N HSQC spectra at NZD:EDTA molar ratios of 1:0 (green), 1:2 (pink), and 1:5 (red). In addition to the coordinating residues highlighted in panel (A), other residues exhibit widespread peak attenuation and chemical-shift perturbations across the titration series. **(C)** Quantification of peak-intensity changes in the ¹H–¹⁵N HSQC spectrum at a protein:EDTA ratio of 1:2. Gray regions denote missing resonance peaks, including prolines (which lack backbone amide protons) and residues with severe peak overlap, preventing confident assignment.

**Figure S4.**
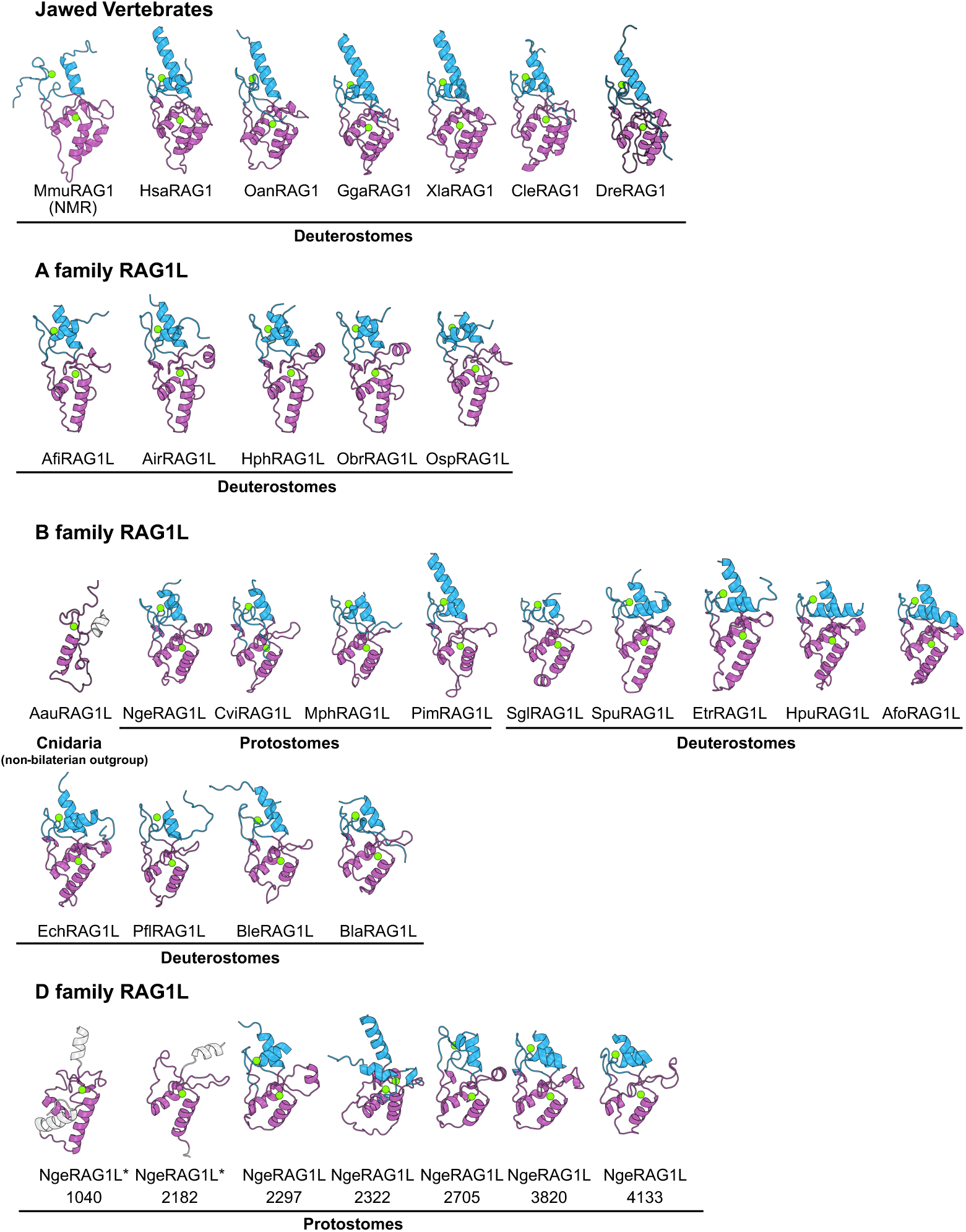
AlphaFold3-predicted RAG1/RAG1L NZD structures across species, related to Figure 3. AlphaFold3 (AF3)-predicted structures of NZD from representative jawed vertebrate RAG1 and invertebrate RAG1L proteins; the mouse RAG1 NZD is shown as the experimentally determined NMR structure. Coloring follows Figure 3A: ZFa in purple, ZFb in light blue, and zinc ions in green. NZD structures are grouped by RAG1L families A, B, and D, and by jawed vertebrate RAG1. Most NZDs contain both ZFa and ZFb zinc-finger modules, whereas AauRAG1L NZD (family B) and Nge1040/Nge2181 RAG1L NZDs (family D) possess only a single ZFa-like module in AF3 predictions and are marked with asterisks (*). Species name abbreviations used in this paper: *Mmu, Mus musculus; Hsa, Homo sapiens; Oan, Ornithorhynchus anatinus; Gga, Gallus gallus; Xla, Xenopus laevis; Cle, Carcharhinus leucas; Dre, Danio rerio; Afi, Amphiura filiformis; Air, Astropecten irregularis; Hph, Hippasteria phrygiana; Obr, Ophioderma brevispina; Osp, Ophyothrix spiculata; Aau, Aurelia aurita; Bbe, Branchiostoma belcheri; Bla, Branchiostoma lanceolatum; Pfl, Ptychodera flava; Spu, Strongylocentrotus purpuratus; Etr, Eucidaris tribuloides; Ech, Evechinus chloroticus; Hpu, Hemicentrotus pulcherrimus; Afo, Asterias forbesi; Cvi, Crassostrea virginica; Sgl, Sphaerechinus granularis; Mph, Modiolus philippinarum; Nge, Notospermus geniculatus; Pim, Pinctada imbricata*.

**Figure S5.**
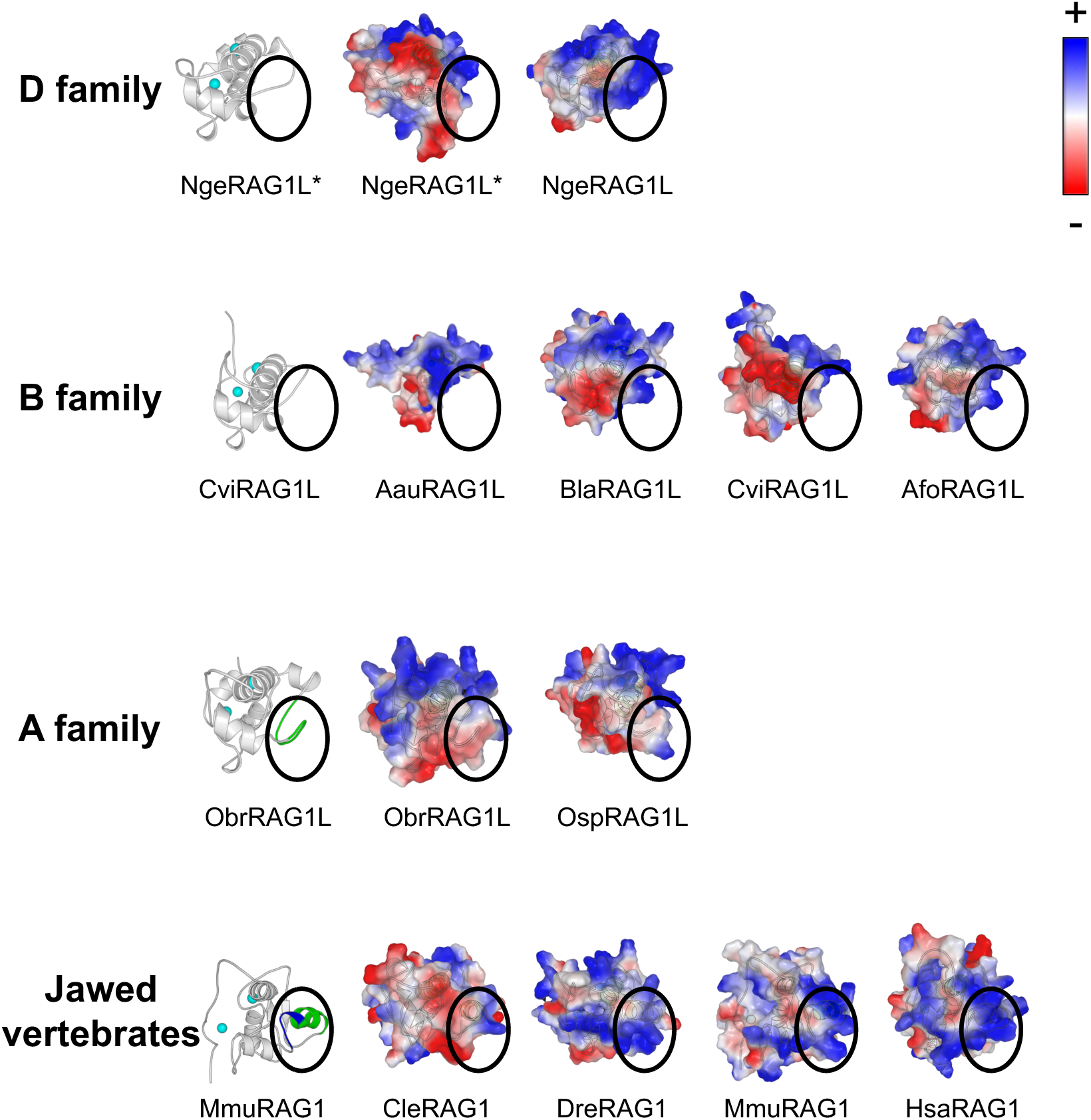
H2 helix and its subsequent basic loop generate a new positively charged surface, related to Figure 3. Electrostatic surface potential variation of the newly formed basic face of NZD across RAG1/RAG1L families. In jawed vertebrates, the H2 helix and subsequent positively charged residues generate a distinct electropositive patch on the bottom surface (outlined with a black dashed circle). *Indicates that this NZD contains only the ZFa module.

**Figure S6.**
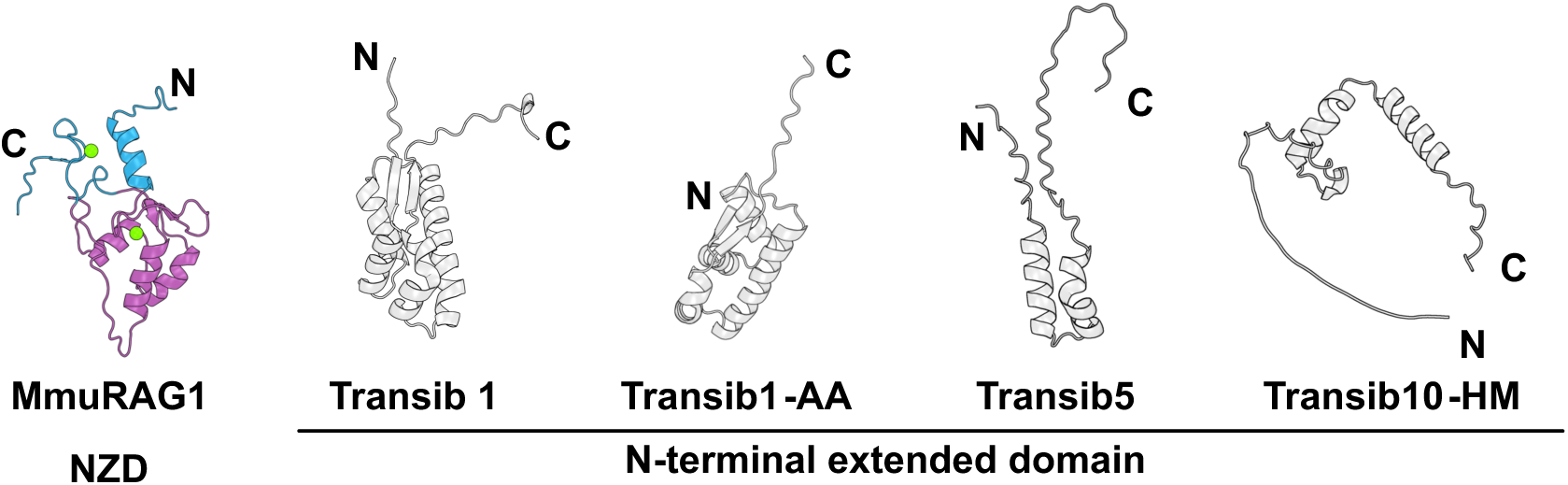
Structure prediction of the N-terminal domains of representative Transib transposases, related to Figure 4. AlphaFold3 predictions of the N-terminal undetermined region from multiple Transib families show structural diversity and distinct differences from the mouse RAG1 NZD.

**Figure S7.**
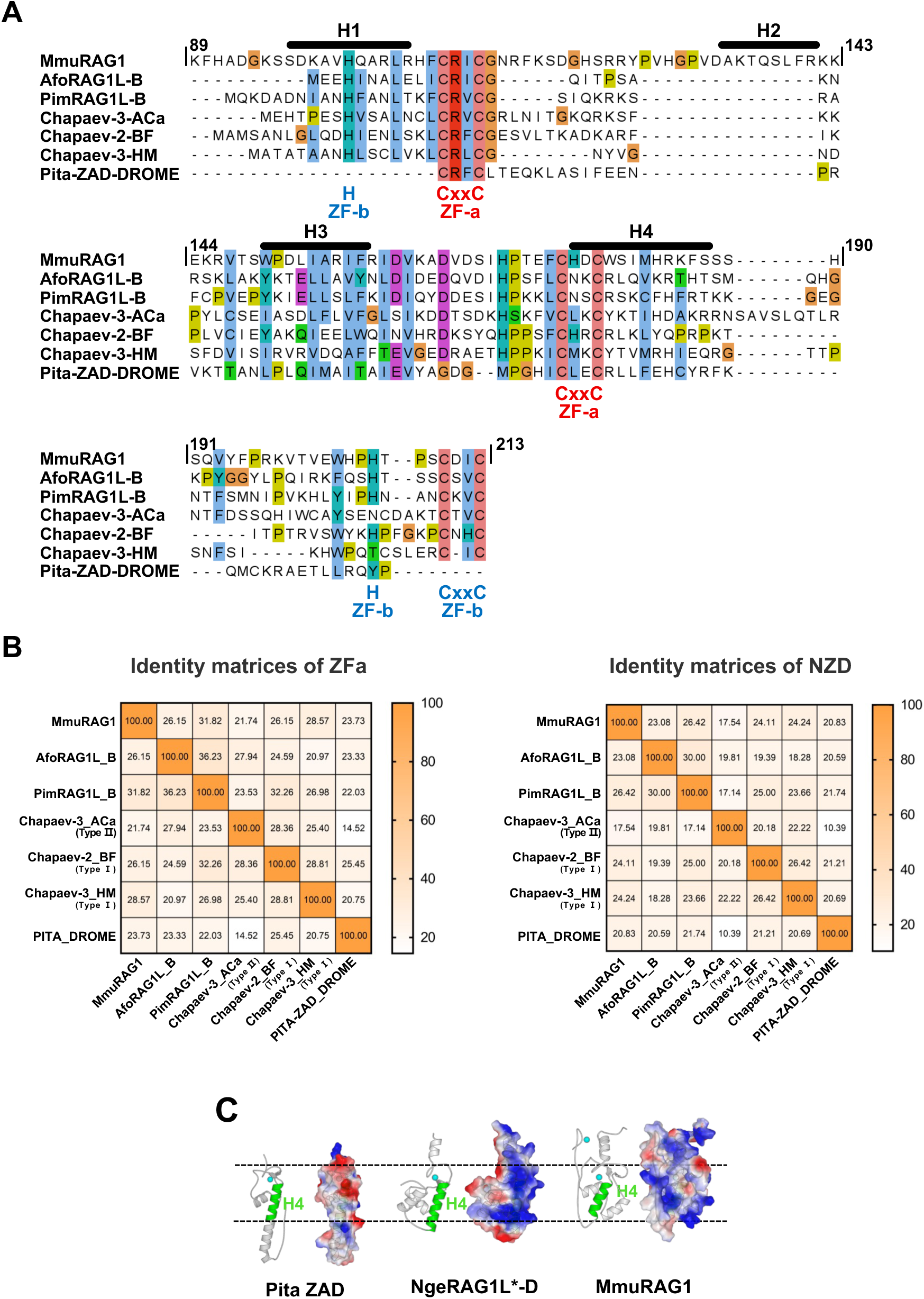
Comparative analysis of ZFa modules within NZDs, and Chapa domains and ZAD, related to Figure 4. **(A)** Multiple sequence alignment of NZDs from mouse RAG1 and selected members of B-family RAG1L, Chapa domains from indicated Chapaev transposases, and the ZAD of Pita. **(B)** Pairwise identity matrices for ZFa (mouse RAG1 110–187) and the intact NZD/Chapa domain among mouse RAG1, selected B-family RAG1L, type I and type II Chapaev (Chapa domains), and Pita ZAD, as shown in (A). Left panel: ZFa only; right panel: intact NZD (RAG1/RAG1L) or Chapa domain (Chapaev). **(C)** Electrostatic surface potential comparison between Pita ZAD and RAG1/RAG1L NZD reveals distinct features in the zinc-finger and H4 regions, with Pita lacking the prominent electropositive patch observed in RAG1/RAG1L.

